# Non-invasive in-blood glucose sensing

**DOI:** 10.1101/2022.11.30.518508

**Authors:** Nasire Uluç, Sarah Glasl, Dominik Jüstel, Miguel A. Pleitez, Vasilis Ntziachristos

**Author notes:** Corresponding authors: Miguel A. Pleitez,; Vasilis Ntziachristos.

## Abstract

Non-invasive glucose monitoring (NIGM) is increasingly considered as an alternative to finger pricking for blood glucose assessment and management of diabetes in insulin-dependent patients, due to the pain, risk of infection, and inadequacy of finger pricking for frequent measurements. Nevertheless, current NIGM techniques do not measure glucose in blood, but rely on indirect bulk measurement of glucose in the interstitial fluid, where glucose is less concentrated, diluted in a generally unknown volume, and appears in a delayed fashion relative to blood glucose, impairing NIGM accuracy. We introduce a new biosensor, termed <u>D</u>epth-gated mid-Infra<u>R</u>ed <u>O</u>ptoacoustic <u>S</u>ensor (DIROS), which offers for the first time non-invasive glucose detection directly in blood, while simultaneously rejecting contributions from the metabolically inactive *stratum corneum* and other superficial skin layers. This unique ability is achieved by time-gating mid-infrared optoacoustic signals to enable glucose readings from depth-selective localization in the microvasculature of the skin. In measurements of mice *in vivo*, DIROS revealed marked accuracy improvement over conventional bulk-tissue glucose measurements. We showcase how skin rejection and signal localization are essential for improving the NIGM accuracy, and discuss key results and how DIROS offers a holistic approach to address limitations of current NIGM methods, with high translation potential.

## Introduction

Blood glucose monitoring remains the main management strategy for prevention of diabetes-related complications. Until recently, glucose measurements relied on electrochemical sensing that required blood extraction via finger pricking, a procedure that is painful, damages tissue, and can cause infection. Intracutaneously implanted electrochemical biosensors integrated on flexible wearable patches have been proposed^1–4^ for minimally invasive glucose monitoring in interstitial fluid or sweat. These methods allow for frequent glucose sampling that improves diabetes management.^5^ However, measurements using implanted biosensors only assess glucose diluted in the interstitial fluid (ISF) upon diffusion from blood capillaries, which decreases the accuracy of readouts^6–8^. In particular, ISF glucose appears in a delayed fashion and at much lower concentrations (up to 10x) compared to clinically-relevant blood glucose. Moreover, glucose concentrations in ISF depend on ISF volume and biochemical environment. Therefore they may be affected by the levels of hydration or pH values. For those reasons, measurements based on implanted electrodes require frequent re-calibration of the implanted sensor using finger prick measurements. In addition, the invasiveness and need for frequent replacement of the measuring electrode carries the risk for skin irritation and microbial infections.

Non-invasive glucose monitoring (NIGM) is heralded as the next frontier in diabetes management, to minimize risk of infections and re-calibration needs over implantable biosensors, while improving the accuracy of blood glucose measurements^9-11^. Beyond improving diabetes management, NIGM could become a pivotal technology for prevention or early detection of diabetes in high-risk populations and as part of a wellness society in need of informed biomedical readings for achieving a healthy lifestyle.

The need for NIGM is underscored by the wealth of methods considered for its implementation. Terahertz (THz) spectroscopy uses the absorption spectrum of glucose (0.1 – 2.5 THz) to assess its concentration in human skin^1–4,12^; however, similarly to implanted biosensors, it is limited to bulk glucose measurements. Moreover terahertz spectroscopy operates with low signal-to-noise ratios, broad absorption bands, and overlapping spectra of glucose with other biomolecules, i.e. parameters that challenge the sensitivity and specificity requirements of glucose monitoring in tissues^13^. With a broader track record, different optical methods have also been considered for NIGM over the past decades^3,13-15^. Raman spectroscopy achieves biomolecular specificity^16^ by resolving specific vibrational spectral signatures of glucose at the fingerprint region of carbohydrates (1300-900 cm^-1^)^17-18^. Nevertheless, the method notoriously suffers from the weak Raman scattering cross-sections of biomolecules, which reduce the detection sensitivity. Based on light absorption rather than scattering, mid-infrared (mid-IR) spectroscopy using optical, optoacoustic (photoacoustic), or photothermal detection has been considered to improve sensitivity at spectral ranges similar to those used in Raman spectroscopy^19-24^. However, conventional mid-IR spectroscopy suffers from difficulties in operating in reflection mode for detection and reproducibility issues associated with measurement contamination from non-glucose-specific absorption of light at superficial skin layers, further rendering calibration methods ineffective^8^.

Critically, as transdermal methods, both Raman and mid-IR optoacoustic spectroscopy perform bulk measurements of glucose in interstitial fluid. Consequently, none of these methods have offered so far a viable alternative to implanted or invasive biosensors for continuous glucose sensing^25-26^.

We present herein a non-invasive method for *in-blood* glucose sensing, to address the limitations of previous methods that operate with bulk glucose measurements in ISF. Termed Depth-gated mid-IR Optoacoustic Sensing (DIROS) the method operates by time-gating optoacoustic signals generated by mid-IR excitation.^27^ We hypothesized that depth-gating would significantly improve the sensitivity and accuracy of glucose sensing based on two key premises:

❶ It would enable the rejection of contributions from the metabolically inactive stratum corneum and overall from the epidermis, since changes in skin humidity, superficial lipids and other molecules are known to contaminate glucose measurements and challenge their reliability and reproducibility.^25-26^
❷ It would allow the detection of signals from volumes rich in microvasculature, i.e. blood-filled volumes. The concentration of glucose in blood is more clinically relevant and, advantageously, much higher than in ISF. In-blood sensing also reports glucose fluctuations in real time, contrary to measurements of ISF glucose that appear in a delayed manner.

Of critical importance in proving these two hypotheses was the depth that could be reached by mid-IR excitation using optoacoustic detection, so that rejection of signals from the epidermis is feasible and subcutaneous microvasculature rich volumes could be interrogated. For this reason, we rigorously examined for the first time the depth achieved by mid-IR optoacoustics, *in vivo*. In particular, we employed a broadband ultrasound detector (bandwidth ∼ 6-36 MHz) to examine the depth achieved and separate tissue layers, with an axial resolution of <25 microns. To further improve the accuracy of the depth investigation, mid-IR measurements were contrasted to congruent microvasculature-sensitive optoacoustic measurements at 532 nm illumination, the latter serving for validation of the depths and structures probed. Then, using the merits of depth-selective optoacoustic detection, we explicitly examined the effects of rejecting signals generated by the epidermis, validating the key hypotheses stated above. In the following, we present the results of the interrogations, the DIROS glucose detection performed in a depth-specific manner over bulk measurements and discuss how DIROS can operate by accessing the capillary-rich layer of the human skin to offer an optimal solution toward non-invasive glucose monitoring for improving diabetes management.

## Results

DIROS was implemented using a common optical path for mid-IR and 532 nm illumination so that mid-IR measurements could be referenced to vascular features that are easily detected in the visible. The optical path (**Fig. 1a**) consisted of a pulsed mid-IR beam (20 ns pulse duration; 2941-909 cm^-1^ / 3.4-11μm spectral range) and a co-aligned 532 nm pulsed beam (5 ns pulse duration); both beams were focused to the surface of tissue (mouse ear) by a broadband reflective objective (see **Methods** for details). Optoacoustic measurements were collected *in vivo* by a focused ultrasound transducer (central frequency of 21.2 MHz with -6dB bandwidth of 72%). For simplicity, the transducer was placed on the opposite side of the tissue measured, establishing a slab geometry. However since the optoacoustic signal is emitted isotopically, operation on the same side of the tissue is also possible. For referencing purposes, we raster scanned the sample under the sensor and generated merged mid-IR/VIS optoacoustic image of tissue (**Fig. 1c**) for use as anatomical references (*see* ***Methods***), in particular images of microvasculature at the 532 nm. Depth selection was implemented by gating the time-dependent optoacoustic signal (**Fig. 1d**) to reject optoacoustic signals generated from the epidermis (**Fig. 1b, EP**), thus minimizing the dependence of the measurement on contributions that do not directly relate to glucose concentration. Optoacoustic signals were processed by the Hilbert transform, so that the results relate to the energy of the signal measured.

**Figure 1:**
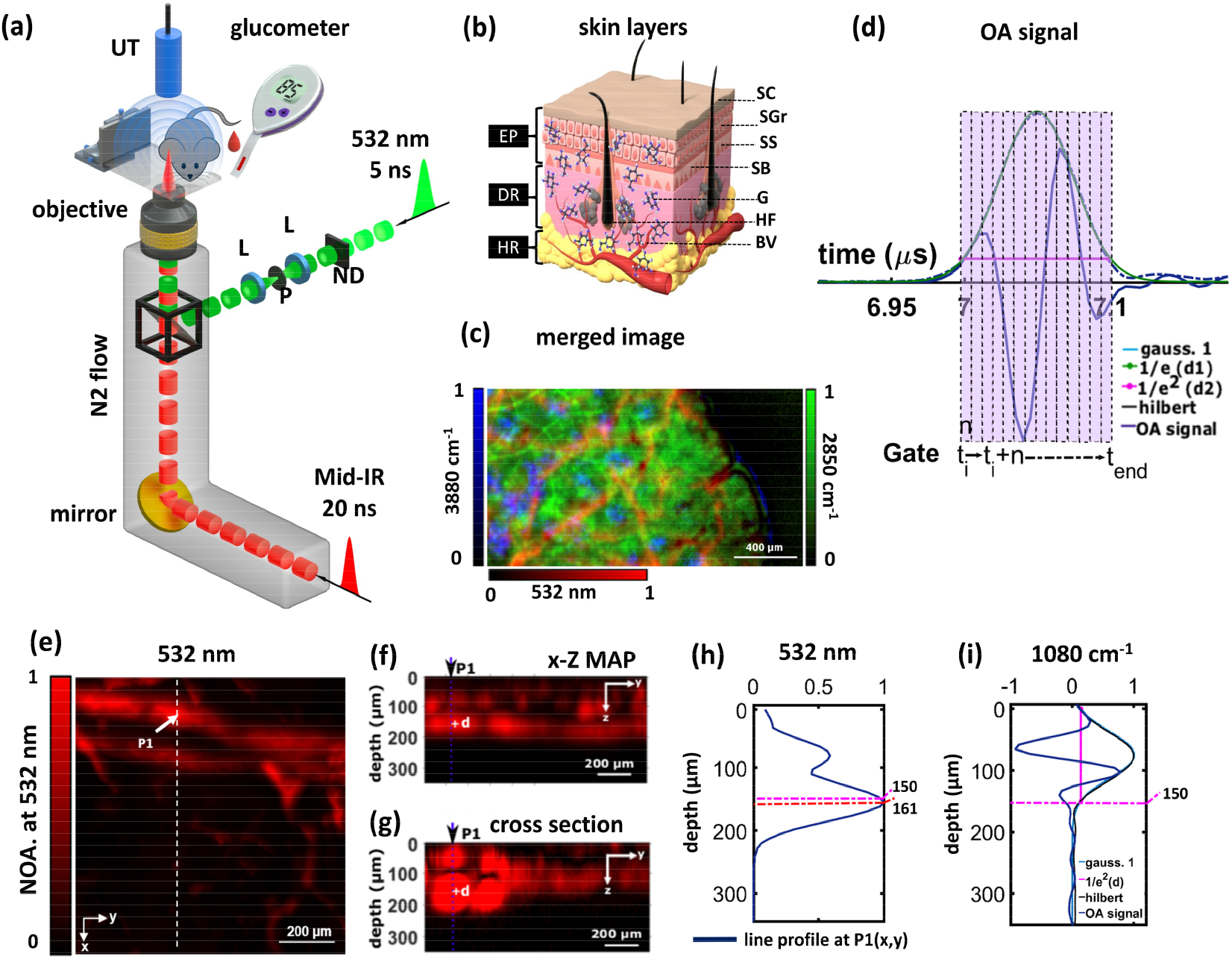
Label-free biomolecular imaging-depth capabilities of *in vivo* visible/mid-IR optoacoustic microscopy. (a) Schematic diagram of the combined mid-IR/VIS *in vivo* optoacoustic microscopy system for image-guided non-invasive glucose monitoring (ND: neutral density filter, L: lens, P: pinhole). (b) Schematic representation of different layers of the mouse skin (EP: epidermis, DR: dermis, HR: hypodermis, SC: stratum corneum, SGr: stratum granulosum, SS: stratum spinosum, SB: stratum basale, SG: sebaceous glands, HF: hair follicles, BV: blood vessels), (c) Merged mid-IR/VIS optoacoustic images, (d) raw optoacoustic signal corresponding to glucose wavenumber (1080 cm^-1^), (e) Representative xy-projected mouse ear image at 532 nm. (f) Maximum amplitude xz-projected image with depth. (g) x-z cross section image, with d (150 μm) representing widths of 1/e^2^ of the Hilbert transform of the optoacoustic transient at 1080 cm^-1^, respectively. (h) Line profile of a blood vessel cross-sectional image (at 532 nm), shown in blue in (f-g). (i) the depth profile of mid-IR with the width of 1/e^2^ of the Hilbert transform of the optoacoustic transient at 1080 cm^-1^, NOS = normalized optoacoustic signal representing widths of 1/e^2^ of Hilbert transform of the optoacoustic transient at 1080 cm^-1^.

The first critical parameter investigated was the depth that can be reached in mid-IR optoacoustics. We previously postulated that, compared to conventional mid-IR optical measurements, mid-IR optoacoustic sensing can interrogate deeper in tissues since it employs ultrasound and not optical detection, i.e. it operates under strong optical attenuation only on the incident but not the collection path^27^. To examine this postulation, we obtained three dimensional micro-vasculature maps from 1×1 mm^2^ scans from mouse ears *in vivo*, using 532 nm excitation (**Fig. 1e-g)**. Maximum amplitude projection (MAP) along the three dimensions allow to observe vascular rich volumes, as exemplified in the images. Then, we plotted the signal profiles at 532 nm (**Fig. 1h**) and 9259 nm (**Fig. 1i**; 1080 cm^-1^) collected from a volume with strong vasculature (P1 position, see below) (see Fig. 1e-g), in order to assess the penetration depth achieved by mid-IR measurements. The P1 position was selected for demonstration purposes, since the capillary present at this position reached the deepest in the volume examined, i.e. a depth of >200 micrometers. The 9259 nm wavelength (1080 cm^-1^ wavenumber) was selected for the mid-IR measurement as a representative wavelength for glucose sensing (see **Fig. 4h**). The penetration depth was defined as the width of 1/e^2^ of a Gaussian curve fitted to the Hilbert transform of the optoacoustic signal at 1080 cm^-1^, as shown in **Fig. 1i** for a measurement that reached a depth of ∼150 μm. The depths reached in all mice are summarized in **Suppl Fig. 1**, showcasing an average depth of ∼140 μm. Such depths are sufficient to measure the capillary-rich layer residing in the epidermal-dermal junction of human skin, not only for mouse measurements. In particular, our own measurements of human skin using raster scan optoacoustic mesoscopy^28^ (**Suppl. Fig. 2**) and independent measurements using optical microscopy^29^ confirm that capillary loops constitute a homogeneous rich-capillary layer that resides at depths of ∼60-80 microns at many sites on the human skin, and is not affected by diabetes progression. DIROS would be then applied to measure from this layer, i.e. the performance demonstrated herein in mice can be transferred to human measurements (see Discussion).

**Figure 2:**
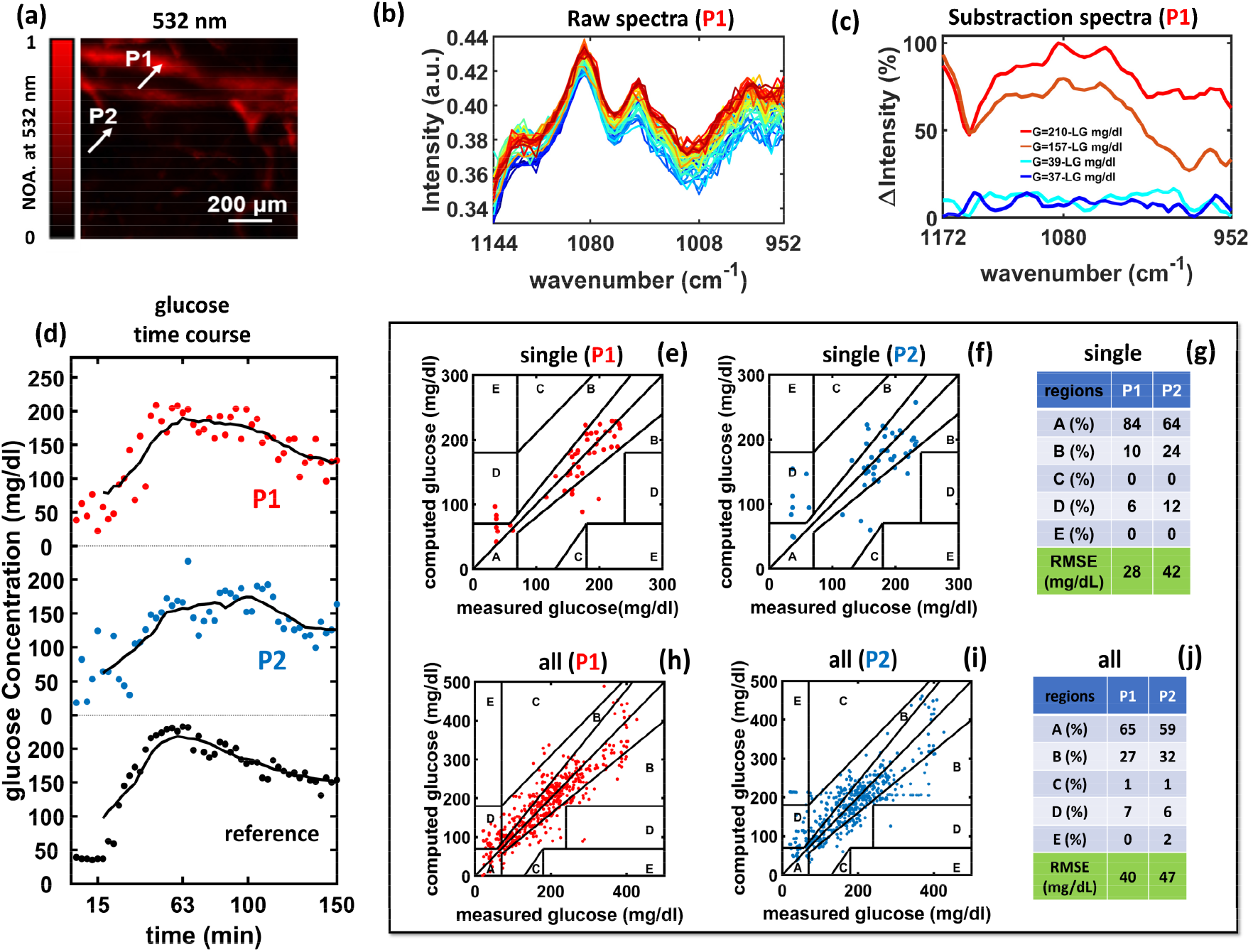
Image-guided non-invasive glucose monitoring with *in vivo* MiROM. (a) Optoacoustic image of a mouse ear at 532 nm, (b) the spectra over time at P1 (c) two spectra corresponding to the lower glucometer value post glucose administration, i.e. at 37 and 39 mg/dL, and two spectra corresponding to the highest values observed in the mouse measurement shown, i.e. at 157 and 210 mg/dL. (d) Time profile of calculated glucose concentrations at P1 and P2 compared to reference blood glucose measurements. (e-f) Clarke error grids showing the correlation between reference and optoacoustic glucose values at P1 (e) and at P2 (f). (g) Tabulation of the distribution of results per region and root mean square errors (RMSECV) by cross-validation for (e-f). (h-j) The Clarke error grids and tabulation of zone distribution for all 10 mice measured in the study for positions P1 and P2.

Having confirmed that the major premise for depth dependent detection from capillary rich layers is feasible, we interrogated the glucose detection performance of DIROS, in relation to in-blood vs. ISF measurements. To achieve this, we performed a glucose tolerance test study in ten mice, based on a 20% glucose solution (2 g/kg of body weight) injected into the abdomen of each mouse. First, a 532 nm absorption map using ∼ 5 μm scanning steps was acquired for each mouse enrolled in the study, to provide a morphological reference of the microvascular distribution in the area under the sensor and to select locations to test in-blood vs. ISF-only measurements. To exemplify performance, we showcase results obtained from the same volume (**Fig. 2a**) used for depth evaluation in Fig. 1. All DIROS scans were performed at two distinct points on the 532 nm maps: a first point (P1) at an area with vasculature presence and a second point (P2) at an area with poor vascularization, (i.e an area representative of measurement in the ISF). Ten baseline spectra (1300 - 900 cm^-1^) were recorded over a time period of 10 min prior to glucose administration and 90 spectra were recorded post-glucose administration over 150 min, continuously alternating the sensor over the positions P1 and P2. Each spectrum was generated as follows: for each of the P1 and P2 locations and wavenumbers scanned, we acquired and added 1000 optoacoustic signals. Each point in the spectrum corresponds to the peak amplitude value of the Hilbert transform of the averaged optoacoustic signal for the selected time gate. Each spectrum consists of measurements at 100 wavenumbers acquired in the 1300 cm^-1^ to 900 cm^-1^ region with a spectral step size of 4 cm^-1^, requiring 1.5 min for acquisition. After one spectrum measurement was completed from one of the two positions selected, the sensor was moved to the other position. For validation purposes, 0.6 μL of blood was obtained from each mouse every 3 min, during the time that a motorized stage moved the sensor from P1 to P2. The blood sample was analyzed by a standard glucometer (see **Methods**). Therefore, each spectrum from the P1 and P2 points corresponds to one reference glucose measurement.

To illustrate the nature of the spectra collected and understand whether the optoacoustic signals respond to different glucose values, we plotted raw spectra (**Fig. 2b**) obtained from position P1 at different time points, hence corresponding to different blood glucose concentration values (for colorcoding see **Suppl. Fig. 3b**). Observation of the spectra showed that the intensity changed as a function of glucose concentration, which was found to be linear, as elaborated in **Fig.4**. To illustrate the spectral change as a function of glucose concentration, we subtracted one baseline spectrum, obtained prior to the administration of glucose, from 4 spectra obtained post glucose administration (**Fig. 2c**): two corresponding to the two lowest glucose concentrations (37 and 39 mg/dL) and two corresponding to the highest concentrations (157 and 210 mg/dL) recorded by the glucometer. These four spectra show that in all cases there is a clear difference in spectrum over baseline that is well above the noise level. The change in intensity observed for the low glucose values is ∼20% of the maximum change in intensity observed in the data set collected, confirming sufficient signal-to-noise ratio for *in vivo* glucose detection at physiological concentrations. A plot of all difference spectra in **Fig.2b** are shown in **Suppl. Fig. 3a**.

To quantitatively investigate the relation between spectral changes and blood glucose concentration, beyond the observation of raw spectra, we employed a multivariate analysis (MVA) method based on the partial least squares regression (PLSR) method. MVA is the typical approach for computing analyte concentrations from spectroscopic glucose sensors^30^ and it considers the structure of the entire spectrum (100 variables) for computing a single glucose value, in the presence of other contributors (metabolites) in tissue. Given a number of spectra (measurements) and ground truth glucose values (obtained from the glucometer), the PLSR describes the spectral data as a linear combination of a new set of spectral components (basis spectra), and identifies the subset of components that is maximally informative of the glucose level. Then, it computes a glucose value based on the particular combination of these spectral components that describes a given spectrum. We applied a leave-one-out cross correlation, whereby each spectrum employed for a glucose measurement was excluded once from the decomposition to basis spectra (see **Methods**) to determine features that represent spectral variation.

Using MVA analysis, we plotted the glucose values obtained from P1 and P2 versus the glucometer values over the time course of a measurement (**Fig. 2d**). While both P1 and P2 tracked the administration of glucose, it is clear that the data from location P1 more closely resembled the blood glucose dynamics recorded by the glucometer. The curves also show a delayed appearance of glucose, when measured at position P2, consistent with the fact that glucose changes in interstitial fluids, represented herein by measurements at P2, appear in a delayed manner compared to the dynamics of blood glucose, represented by measurements at P1. Time course control experiments injecting phosphate buffer saline (PBS) were also performed in 3 mice. The results showed a minimum baseline increase that was virtually constant throughout the time course of the measurement (**Suppl. Fig. 4**), supporting that the signals in **Fig. 2d** are due to the glucose injection.

To study the accuracy of the glucose measurement, in relation to the two measurement locations, we plotted the Clarke Error Grid (CEG) analyses for locations P1 and P2, for the mouse shown in Fig. 2a (**Fig. 2e-g**) and for the entire data set collected from all mice (**Fig. 2h-j**). The CEG is divided into five regions (A-E), representing degrees of accuracy of glucose estimations. Values falling into different zones have various levels of inaccuracy, with values within zone A being the most accurate (within 20% of the reference measurement), while those in zones D and E represent erroneous readings^31^. Visually, the results for P1 appear less scattered and better confined in the A area compared to the measurements at P2. Correspondingly the root-mean square error cross-validation (RMSECV) value between the measurements at P1 and P2 and the reference glucometer values were found to be 28 mg/dL vs. 42 mg/dL for the single mouse and 40 mg/dL vs. 47 mg/dL for the entire cohort.

The glucose measurement results shown in **Fig. 2** were obtained only by selecting areas with and without vasculature but without depth selectivity, confirming the hypothesis that measurements from blood-rich volumes are more accurate than measurements in ISF. The next critical step, and the key point of the development of the DIROS sensor, was to examine whether depth-selection could further improve the performance beyond the capabilities of current sensors. Time gating avoids bulk measurements and can localize readings from blood-rich layers or volumes that lie under the epidermis. Therefore, this approach avoids non-specific contributions from the epidermis and bulk ISF measurements, offering measurements that can be labelled as in-blood. We note that while the work herein is guided by images, it can be applied *in vivo* without imaging, as elaborated in the discussion, by targeting the epidermal-dermal junction layer that is rich is blood-filled capillaries across the skin in animals and humans (see **Suppl Fig. 2**).

Glucose concentrations were computed with and without gate-selection at points P1 and P2. Results (**Fig. 3**) are showcased using a different mouse than the one shown in **Fig. 1-2** to additionally illustrate the diversity seen in the collected vascular maps. Similar to the analysis in Fig. 2, we selected two measurement locations: one with higher (P1) and one with lower (P2) microvascular density. However, here we applied a time-gate algorithm (see **Methods**) that was optimized so that the OA signal was sectioned to obtain spectra at time gates (depths) that minimized the error between the DIROS measurements (i.e. the Hilbert transform of the optoacoustic signal) and the reference glucose measurements. Different layers correlated differently to the measured glucose values, confirming that DIROS performance varies with depth. A layer at a depth of 97.5 ± 20 micrometers gave an optimal error minimization for all mice studied and was therefore selected as the gate for all mice and all measurements. A first insight into the effects of time gating is seen in **Fig 3b**, which compares glucose values at different gates, i.e. at different skin layers (depths), to the reference measurements and visually shows that the selected time-gate provides the best match. It can be observed that superficial measurements correspond to bulk measurements from the stratum corneum and top of the epidermis, similar to the measurements performed by other sensors, and show the worse match to the glucometer values, offering a first validation of the main DIROS hypothesis that depth selection can improve accuracy. We computed the Pearson correlation coefficient between DIROS measurements and glucometer values to quantify the match between the two techniques. We found a Pearson correlation coefficient of *R* =0.92 for measurements at a depth of 97.5 μm, but lower correlation coefficients of *R* =0. 80 and *R*=0.72 as the gate was moved toward the skin surface.

**Figure 3.**
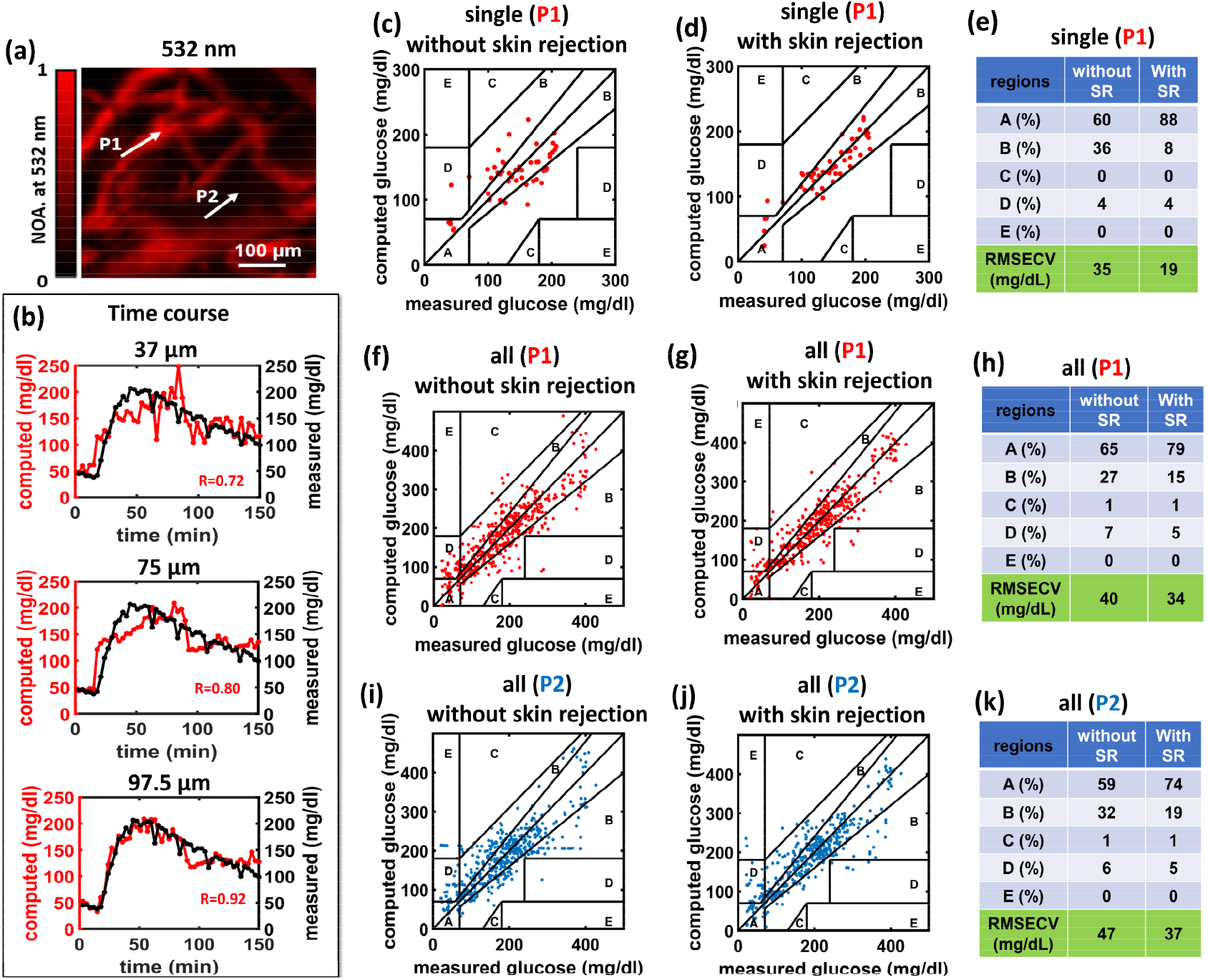
Depth-selective non-invasive glucose monitoring with time-gated optoacoustic sensing. (a) Optoacoustic micrographs at 532 nm (b) glucose time course of blood glucose variation at different depths (37, 75, 97.5 μm) with reference glucose values. A Clarke error grids for a representative experiment in a single mouse when measuring directly at P1: (c) without skin rejection (SR) and (d) with SR. (e) Table comparing the distribution of results per region, and average root mean square error of cross-validation (RMSECV) for P1 with and without SR. A Clarke error grid for 10 experiments when measuring directly at P1: (f) without SR and (g) with SR. (h) Table comparing the distribution of results per region, and average root mean square error of cross-validation (RMSECV) for SR and without SR measurements for P1. A Clarke error grid for 10 experiments when measuring directly at P2: (i) without SR and (j) with SR. (k) Table comparing the distribution of results per region, and average root mean square error of cross-validation (RMSECV) for SR and without SR measurements for P2.

To further validate the effect of depth selection, we plotted the Clarke Error Grid (CEG) with and without gate selection (**Fig. 3c-h**) and show up to **∼** 2-fold sensitivity improvement when using the optimal gate (**Fig. 3e**). The results from the single mouse showcase that measurements from microvascular-rich volumes with depth selectivity **(Fig. 3d)** yielded higher accuracy (88% of the points in zone A) compared to measurements obtained without skin rejection **(Fig. 3c;** only 60% of the points in zone A). When comparing the results from all mice, 79% of the measurement points fell in zone A of the CEG for the P1 location using skin-rejection, whereas only 65% of the measurement points fell in zone A without the time gate. Therefore, the most sensitive performance was achieved for measurements obtained from the P1 position after applying a time gate. Overall, the RMSEs for the entire cohort of mice improved from 47 mg/dL for bulk ISF measurements (P2; Fig. 3k) to 34 mg/dL for measurements of blood-rich volumes with depth selection (P1; Fig. 3h).

To better elucidate the differences in glucose measurements at different time gates (**Fig. 4a**), we plotted the spectra collected from a superficial layer (**Fig. 4b** @37μm) and a deeper layer (**Fig. 4c** @97.5 μm) from location P1 at different time points, i.e. different glucose concentrations. The spectra recorded from the deeper layer show increasing intensities as glucose concentrations increase (for color coding see **Suppl. Fig. 3c**). Furthermore, it is visually evident that the changes in the deeper layer are more prominent than in the superficial layer. Moreover, while spectral changes due to glucose are observable in the superficial layers, water contributes more to the spectral shape within the superficial layers than in the deeper layers, where the spectra more closely resemble that of glucose (see **Fig. 4h**).

**Figure 4.**
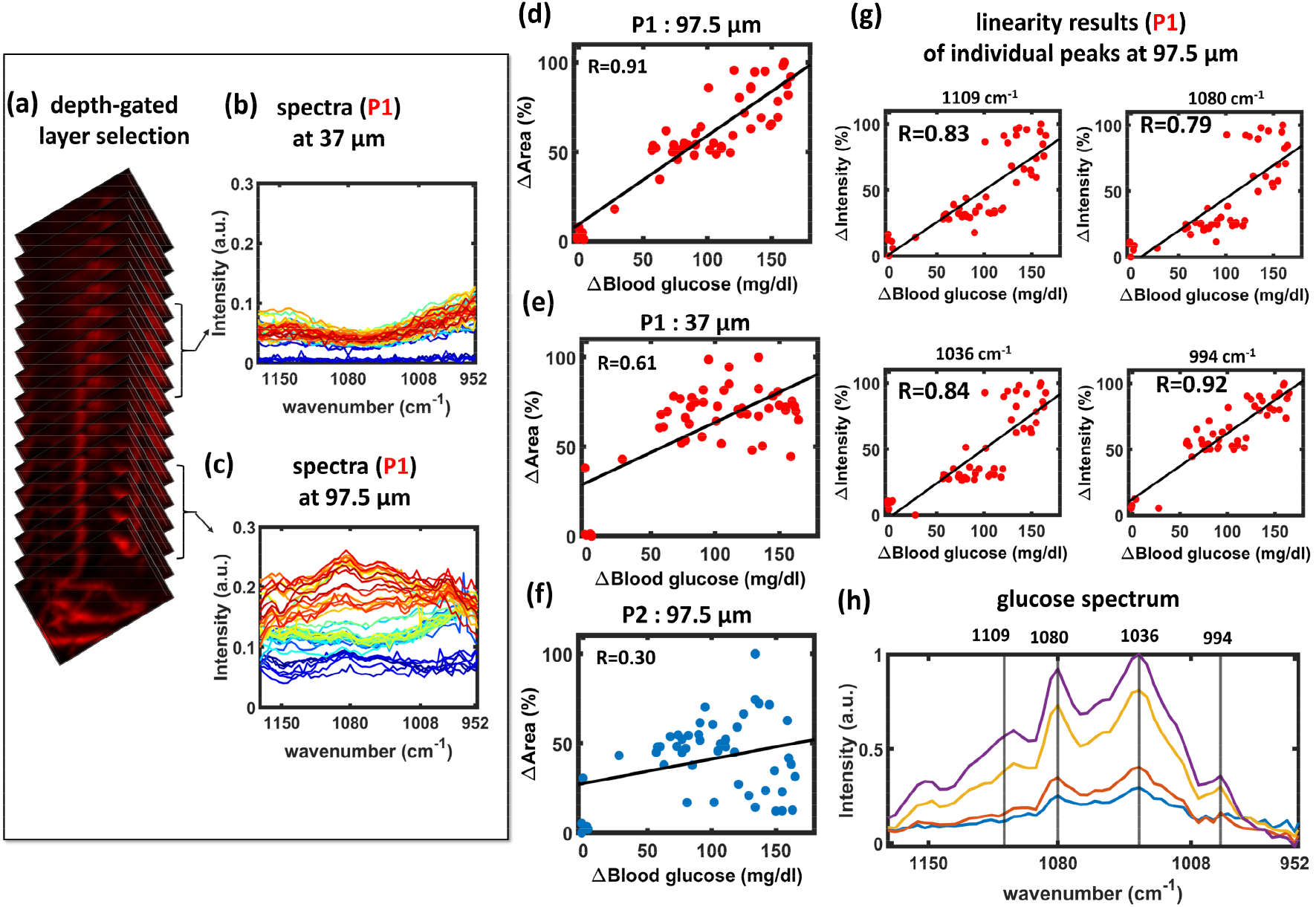
Different time-gates measure different spectral compositions at different skin layers. (**a**) A representation of different skin slices selected by time-gating the optoacoustic signals and corresponding spectra for P1 at depths 37 μm (**b**) and 97.5 μm (**c**). Plot of the change of the area-under the curve of the subtracted spectra as a function of reference glucose values determined by the glucometer for the 97.5 μm layer at P1 (**d**) the 37 μm layer at P1 (**e**), and the 97.5 μm layer at P2 (**f**). Correlation of intensity changes as a function of reference glucose values determined by the glucometer for different peaks in the spectrum collected, in particular corresponding to the 1109, 1080, 1036 and 994 cm^-1^ wavenumbers. (h) Glucose spectrum measured by DIROS in water solution.

To study the linearity of the spectra, we plotted the area under the curve versus glucose concentration for the deep and superficial layers at position P1 (**Fig. 4d-e**) and for the deeper layer at position P2 (**Fig. 4f**). We observed approximate linear correlations at all locations; however, the best correlation (R=0.91) was obtained for the deeper layer at position P1, which is closer to vasculature and rejects contributions from the skin. Measurements at the more superficial layer at position P1 gave a correlation coefficient of R=0.61, whereas measurements from the deeper layer at the poorly vascularized position P2 exhibited the worse correlation (R=0.30).

While these area plots are useful in understanding the energy signal of the entire measurement, we were also interested in investigating whether individual wavenumbers (wavelengths) would suffice for glucose prediction. Therefore, we plotted the intensities of four different wavenumbers corresponding to peaks in the glucose spectrum (**Fig. 4h**), obtained from the 97.5 μm layer of P1, as a function of glucose concentration (**Fig. 4g**). Individual wavenumbers also showed good correlation with the measured glucose values with the peak at 994 cm^-1^ demonstrating the highest correlation of R=0.92.

## Discussion

We have demonstrated position and depth-selective glucose sensing *in vivo* by combining visible optoacoustic microscopy with mid-IR optoacoustic microscopy/spectroscopy for direct glucose-in-blood sensing. First, we demonstrated that mid-IR optoacoustic sensing can reach an average depth of ∼140 micrometers, i.e. a depth capable of reaching skin layers rich in vasculature, in particular the epidermal – dermal junction, as illustrated in Suppl. **Fig. 2**. Then we interrogated whether detection in blood-rich volumes with and without skin rejection improves the sensitivity and detection accuracy. Measurements over time (**Fig. 2d**) and at different depths (**Fig. 3b**) consistently demonstrated the major DIROS hypothesis; namely, that measurements in blood-vessel-rich volumes with skin detection improve the performance over current sensing approaches based on bulk glucose measurement in ISF.

To the best of our knowledge, this is the first report of a sensor capable of in-blood non-invasive glucose measurements. By rejecting skin contributions, DIROS is shown to improve upon one of the major challenges of other optical sensors, i.e. the contamination of the measurement due to mid-IR contributions from parameters in the skin epithelium, including sweat, levels of hydration or lipids. Besides challenging absolute quantification, these parameters are known to contribute to a highly heterogeneous skin appearance in the mid-IR, making optical NIGM measurements unreliable.^22^ Therefore, DIROS improves upon previous NIGM technologies by combining the high glucose OA signal generated by mid-IR absorption with minimization of other strong absorbers that challenge the accuracy and repeatability of bulk measurements.^25-26^

A particular feature of the study herein has been the co-registered use of 532 nm optoacoustic measurements to produce reference frames of accurate microvascular characterization. Moreover, we opted for broadband ultrasound detection combined with focused illumination to allow for morphologic and metabolic investigations of skin heterogeneity and of the DIROS main hypotheses with high positional certainty afforded by the system’s high spatial (6 micrometers) and axial (25 micrometers) resolution. The result is a sensor that offers to the best of our knowledge the most accurate in-blood non-invasive glucose detection today.

We presented both observations of raw and multivariate data analysis. Our aim was to show that the sensitivity of DIROS suffices for detecting clear changes in the raw data due to glucose variations, rather than indirect methods based on statistical observations as is typical in the literature. **Fig. 2b** in particular shows clear spectral variations as a function of glucose concentration, demonstrating for the first time in mid-IR glucose investigations *in vivo* that even individual points in each spectrum can vary with glucose concentration in a linear fashion (Fig. 2d). This rather subtle point is critical in understanding the measurements in the forward sense and not only as the output of MVA, which generally acts as a “black box” in data analytics. Similar concerns were recently raised for Raman spectra, with one investigation focusing on a similar demonstration to that shown here in **Fig. 2b-c**, i.e. showing for the first time raw Raman spectra varying as a function of glucose values in vivo^18^. However, a preliminary glimpse into the sensitivity differences between Raman and DIROS suggests that the Raman spectra showed prominent spectral changes for glucose changes in the 256 – 456 mg/dL range, whereas DIROS raw spectra herein demonstrated differences for glucose concentration changes as low as 28 mg/dL.

We further demonstrated direct point-to-point comparison of DIROS measurements against the reference glucometer measurements, not only Clarke Error Plots, offering a more direct observation of results. In particular we observed (**Fig. 2d**) that in-blood measurements, i.e. measurements in capillary rich volumes, offered a more precise time course of blood glucose variation than measurements in ISF, although the overall accuracy of measurements at ISF was significantly compromised compared to in-blood measurements. Likewise, **Fig. 3b** provides insight into the potential contamination caused by skin heterogeneity since employing skin rejection when processing the measurement significantly lowered the error in the computed of glucose concentration compared to when skin was included. At the optimal gate of 97.5 micrometers, there is clearly a much closer point-to-point match between the DIROS and glucometer values, corroborating the main premise of our depth-selective interrogations.

Certain limitations exist in the study. The measurements are obtained from mice and not humans, due to the unfortunate current situation in Germany in regard to regulation of experimental arrangements, in particular in regard to what appears to be an erroneous interpretation of the new Medical Device Regulation introduced by the European Union in May of 2021. While the EU Medical Device Regulation is clearly aimed at commercial developments regarding the placement of medical products in the market or in patient service, German authorities assume this regulation to also apply to research investigations, significantly challenging translation activity. If the situation resolves in the future, an immediate next step would be to repeat this study in humans. While mouse skin differs from human skin, the depths reached herein offer a very promising outlook for reaching the epidermal-dermal junction of human skin as well. Therefore we expect with very high probability these results to also be confirmed in humans. A second challenge herein was that we did not monitor for sensor intensity fluctuations; therefore our results also comprise background and system fluctuations. We partially compensated for this instability by employing a high number of averages and collected spectral points. We expect that in a second sensor generation, we can implement a reference arm to minimize signal variation issues and reduce the number acquisition points needed collection time and further improving the detection sensitivity.

DIROS could be extended beyond glucose measurements to other metabolites, such as lactate and lipids. This could allow, for instance, to develop a continuous metabolic sensing system to alert of deviations from healthy metabolic parameters. In summary, the method presented here is a powerful new tool for precise determination of clinically relevant blood-glucose levels that could pave the way for significant advances in diabetes management.

## Methods

### Combined Visible and Mid-infrared optoacoustic microscopy

A pulsed quantum cascade laser (QCL) (MIRcat, Daylight Solutions), with a tuning range from 3.4 μm to 11 μm, 20 ns duration, and a repetition rate of 100 kHz was used as the optoacoustic excitation source. Additionally, a 3 ns laser beam at 532 nm (Cobalt) was integrated with a flip-mirror sharing the same optical path of the QCL.(**Fig. 1a**) Both, visible and mid-infrared output laser beams were focused to the sample by a 36X reflective objective. Optoacoustic signals from the sample were detected with an ultrasonic transducer with a central frequency of 20 MHz. To evaluate the co-registration accuracy between the two systems, we obtained carbon tape images at 532 nm, the wavelength employed to enable visualization of hemoglobin-based contrast, and at three specific wavenumbers in the mid-IR range corresponding to glucose, lipid, and protein detection in the skin (1085, 2850, 1587 cm^-1^, respectively; **Suppl. Fig. 5a-c**). Comparison of the line profiles through the image center along the x-axis and the y-axis (**Suppl. Fig. 5d**) showed excellent agreement between all images (**Suppl. Fig. 5e-f**). The merged visible and mid-IR optoacoustic image (**Suppl. Fig. 5i**) revealed slight differences in the spatial localization between the two images, calculated by using 10 μm line profiles along the x- and y-axes (**Suppl. Fig. 5j-k)**. This slight difference was taken as a reference when selecting the blood vessels, and because the vessel diameter for selective localization of glucose monitoring was greater than 10 μm, the selective localization was confined to the inside of the vessels.

### Glucose tolerance tests and in vivo mid-infrared optoacoustic spectroscopy

For location-selective non-invasive glucose monitoring *in vivo*, we first used the visible laser integrated into our MiROM system to localize vascular-rich regions, and the image of a mouse ear was acquired at 532 nm. Images of mouse ear tissue were then acquired at 2850 cm^-1^ with the MiROM system to visualize skin heterogeneity (see **Fig. 1c**). The acquired signals at 532 nm and 2850 cm^-1^ were averaged over 50 consecutive signal cycles. Using these images, we selected two different locations (P1 and P2) to test the correlation between MiROM spectral changes (in the range from 1300 cm^-1^ to 900 cm^-1^) and blood glucose concentration. For this sake, glucose tolerance tests were performed in ten different mice at the two selected locations (P1 and P2); 5 baseline spectra were simultaneously acquired over 10 minutes before glucose injection and 45 spectra were collected for 150 minutes after glucose injection. For each *in vivo* mid-IR spectra, we measured a reference blood glucose value with a glucometer (CONTOUR®) to correlate spectral changes and blood glucose concentration. For each glucose tolerance test, a total of 50 blood glucose reference values and 50 *in vivo* MiROM spectra (per measurement point) were obtained.

### Multivariate analysis

The collected spectra were constructed by taking the maximum intensity of the Hilbert transform applied to the retrieved mid-IR optoacoustic transients. Principal Component Analysis (PCA) was applied to the series of OA spectra collected for each glucose tolerance tests (i.e., 50 spectra per test) to determine their common features. Because the size of spectrum data was smaller than the parameter of independent variables at the wavenumber, a Partial Least Square Regression (PLSR) algorithm and cross-validation were used to calculate glucose concentration. The algorithm enabled the rotation of the coordinate system of the data space and the generation of new components; namely, a latent variable. The algorithm thus maximizes variance and correlation between the variables coming from the measured spectrum data and reference glucose concentrations. A PLSR model was constructed after preprocessing through meanscale, and a leave-one-out cross-validation was performed for each glucose tolerance test (GTT) to obtain the root mean square error of cross-validation (RMSECV). For the PLSR analysis, Matlab (Matlab 2019a, Mathworks, Inc. Natick, MA, USA) and PLS (PLS_Toolbox 8.9.2, Eigenvector Research Inc., Manson, Wash., USA) were employed.

### Skin-sectioning depth-selective glucose sensing

To avoid anatomical structures in skin areas with low glucose content for more precise glucose-in-blood detection, a specific time window of optoacoustic signals was used in order to interrogate deeper seated vessels. The width of the time window (w) was selected to be 7.5 μm in the range of the width of 1/e^2^ of the Hilbert transform of the optoacoustic transient, representing the achievable depth at the corresponding wavenumber for glucose detection of OA spectra at two locations (at P1 and P2). For each w, the window was shifted in 7.5 μm steps, and for each position of the window, a spectrum was generated for the GTT. The PLSR model was constructed for each spectrum acquired by time-gated signals, and a leave-one-out cross-validation was performed for the spectral information corresponding to certain depth layers. The root mean square error of cross-validation (RMSECV) between reference and theglucose values in the specific window was calculated. This process of providing spectral information along the skin depth was used as a skin-rejection window to calculate glucose concentrations only from deeper seated vessels.

### Sample preparation and experimental protocol for in vivo glucose monitoring

All mouse experiments were performed according to the guidelines of the committee on Animal Health Care of Upper Bavaria, Germany. The mice were maintained in an individually ventilated cage system (Tecniplast, Germany) at 22 °C ambient temperature, a relative humidity of ∼50% and a regular 12 hours day/night cycle, in our specific-pathogen-free (SPF) mouse facility at the Center for Translational Cancer Research of the Technical University of Munich (TranslaTUM).

Female Athymic nude-Foxn1^nu^ mice (Envigo, Germany) were selected for the glucose tolerance tests. During all the measurements, the mice were anesthetized with 1,6% Isoflurane (cp-pharma, Germany) and 8l pm oxygen as carrier gas. The mouse heart rate, body temperature, and the SpO2 were controlled by a monitoring device (Physio suite, Kent Scientific, Torrington, USA). The imaging of all the mice was performed on the left ear.

After acquiring the baseline data, 2g/kg (body weight) glucose (Braun, 20% Glucose) was injected into the mouse intraperitoneally. For reference glucose measurements, glucose in the blood was measured in parallel with glucometer measurements using test strips (Contour next, Ascensia Diabetes care GmbH, Germany). The blood was extracted from the caudal vein, and the mice were sacrificed immediately after imaging.

## Acknowledgements

We thank Dr. Andriy Chmyrov for useful discussions and system automation and Robert J. Wilson and Sergey Sulima for editing. The research leading to these results has received funding under the European Union’s Horizon 2020 and Horizon Europe research and innovation programme under grant agreement No 862811 (RSENSE) and No 101058111 (Glumon), from the European Research Council (ERC) under grant agreement No 694968 (PREMSOT) and from the Deutsche Forschungsgemeinschaft (DFG) as part of the CRC 1123 (Z1) and from the DZHK (German Centre for Cardiovascular Research).

## Supplementary Figures

**Supplementary Figure 1:**
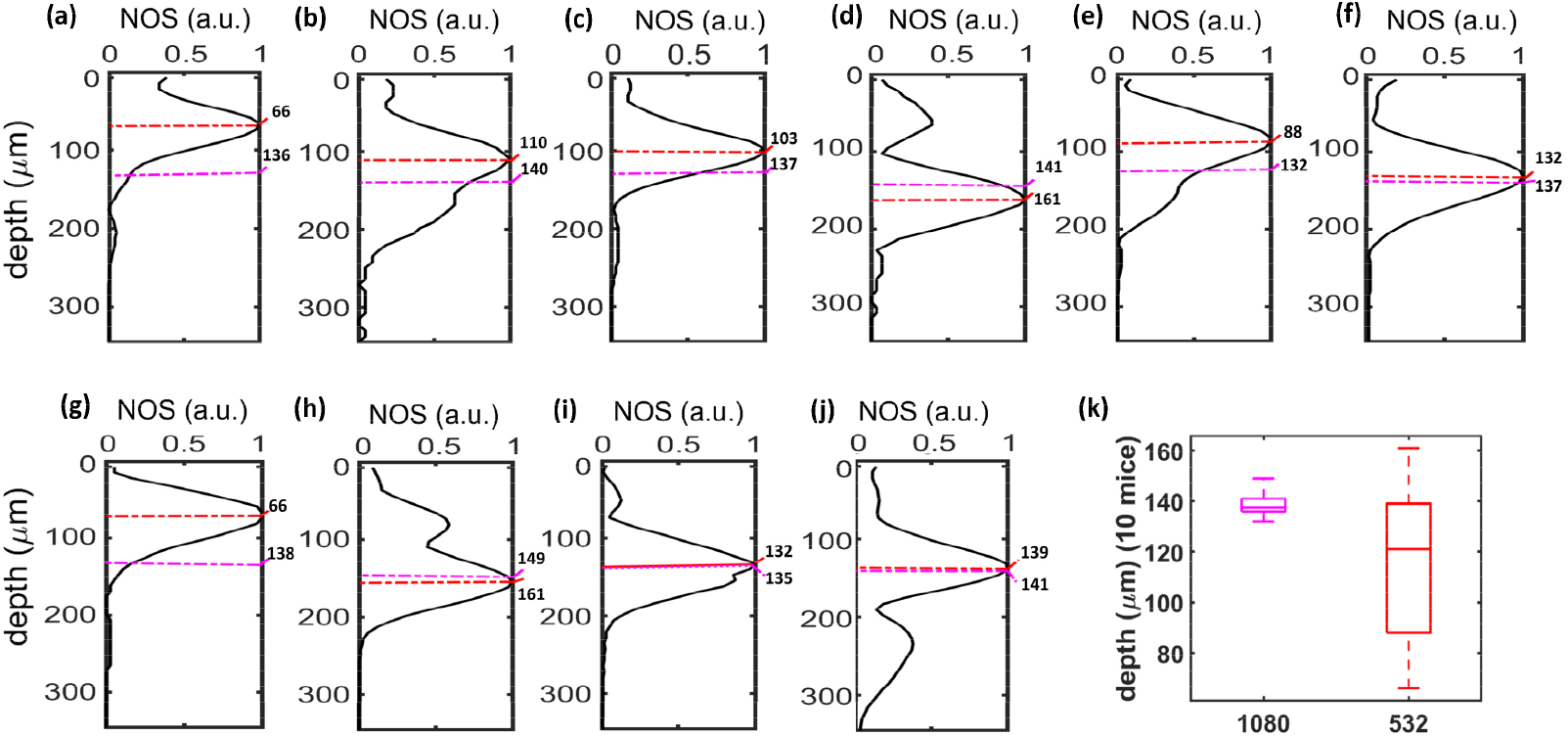
Comparative MiROM/VIS-OA depth analyses at 1080 cm^-1^ and 532 nm for ten experiments. **(a-j)** Line profiles of a blood vessel’s x-z/y-z cross-section images at P1 for the 10 mice studied, with widths of 1/e^2^ of Hilbert transform of the optoacoustic transient at 1080 cm^-1^, and, the maximum intensity position of blood vessel in the depth direction, shown as magenta and red dotted lines, respectively, (k) comparison the depth seated blood vessels and the depth that can be reached in mid-IR optoacoustics sensor for 10 mice, NOAS = normalized optoacoustic signal.

**Supplementary Figure 2:**
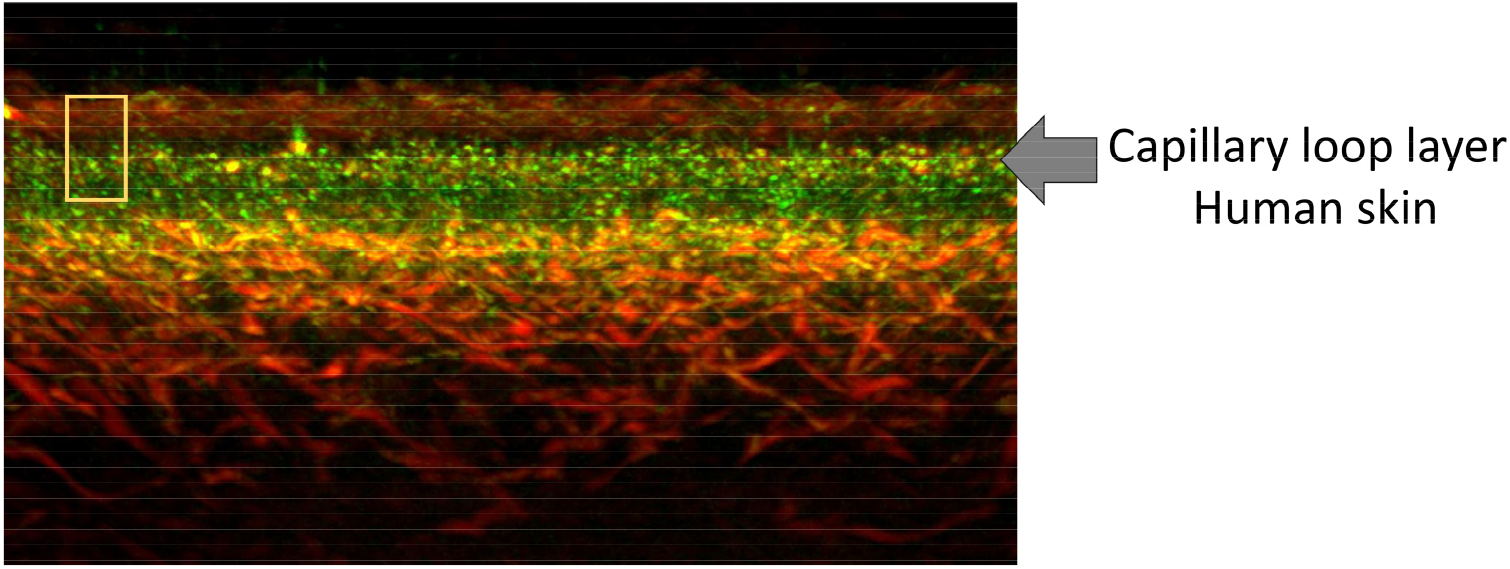
Cross-sectional optoacoustic image from the human skin. The arrow points to the epidermal-dermal junction, which is a superficial layer rich in micro-vasculature. Green color marks the capillary loops in this layer. The yellow box indicates the depth scanned by DIROS, according to the findings of Fig. 1. Image is reproduced From Aguirre J., et. al. Nature Biomedical Engineering 2017.

**Supplementary Figure 3:**
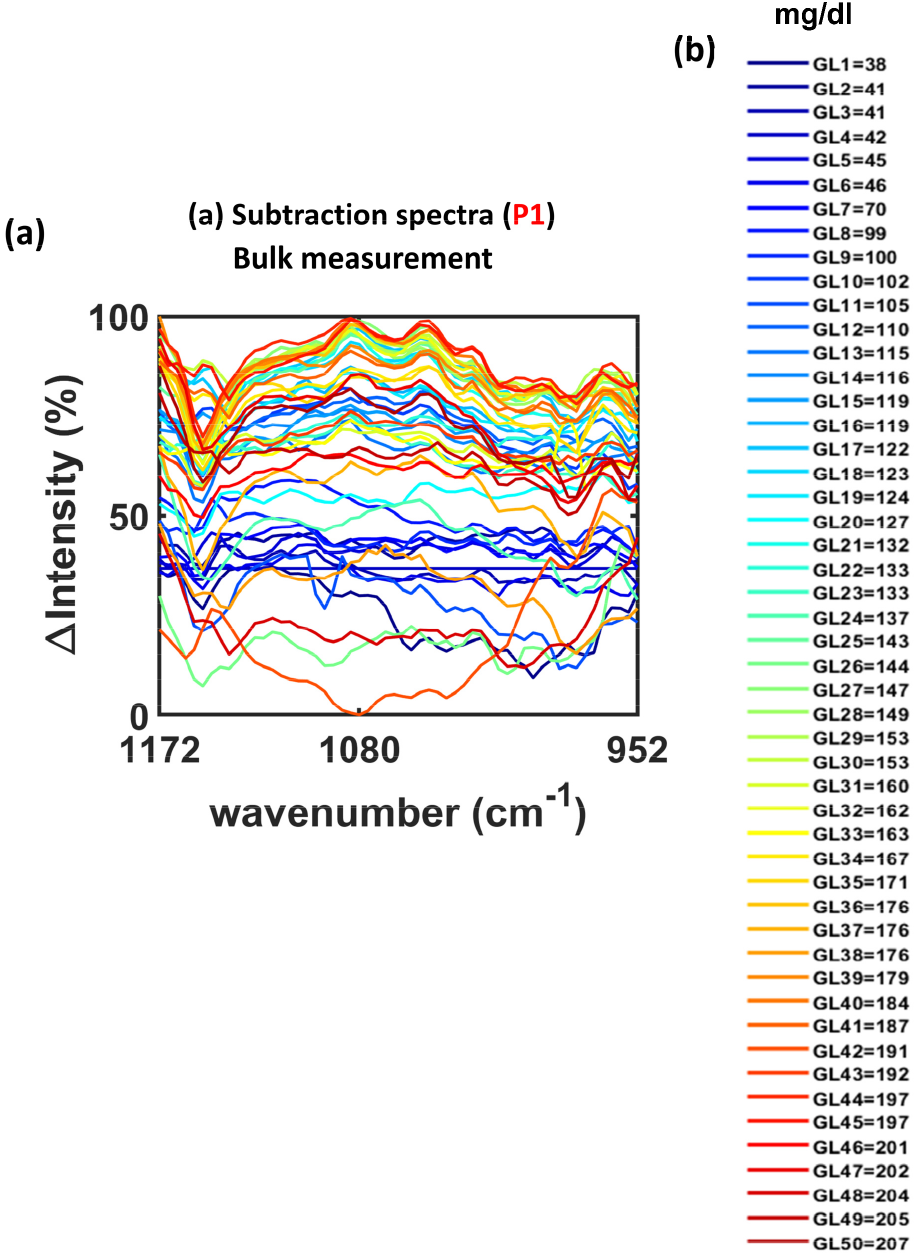
Difference spectra generated as a subtraction of a baseline spectrum (obtained prior to glucose administration) from 50 spectra obtained at different time-points post glucose administration, and hence different glucose values – color-coded in (b). The curves show how bulk measurements are biased toward surface-weighted spectra, especially at the lower glucose concentration vs. depth-selected detection, shown in Fig.4.

**Supplementary Figure 4 :**
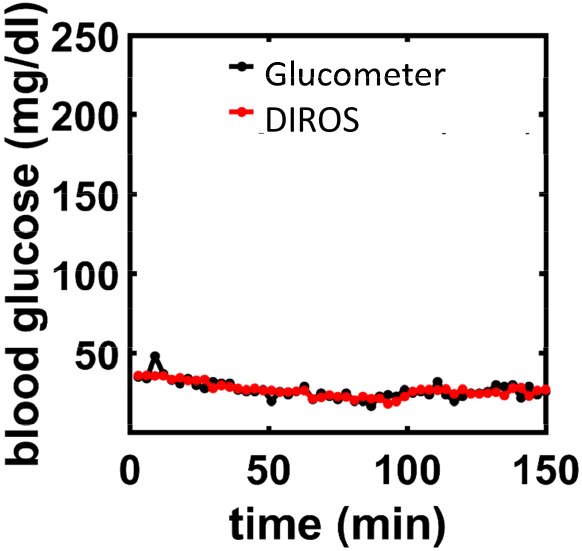
Time course of glucometer and DIROS measurements after injection of PBS in a mouse.

**Supplementary Figure 5:**
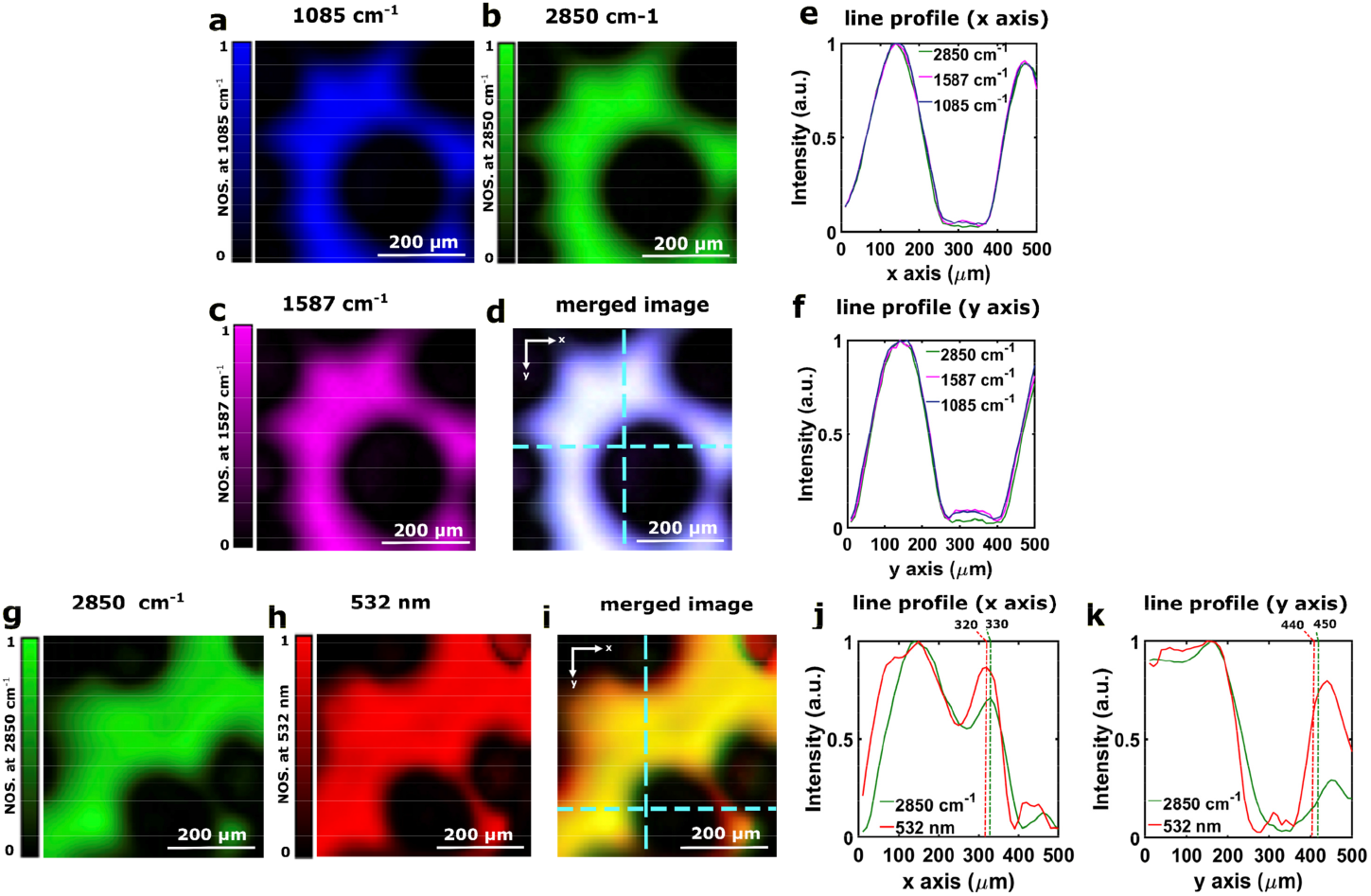
Co-registration accuracy test for the combined VIS-OA/MiROM system. Micrograph of carbon tape at **(a)** 1085 cm^-1^, **(b)** 2850 cm^-1^, **(c)** 1587 cm^-1^. **(d)** Merged image of (a-c). Intensity line profile at the dashed line in (d) along the **(e)** x-axis, **(f)** y-axis. Micrograph of carbon tape at **(g)** 2850 cm^-1^ and **(h)** 532 nm. **(i)** Merged image of (g-h). Intensity line profiles at the dashed line in (i) along the **(j)** x-axis and **(k)** y-axis.

